# Integrated Multi-Omics Characterization of CDK4/hTERT-Immortalized Ovine Satellite Cells for Cultivated Meat Applications

**DOI:** 10.1101/2025.10.23.684220

**Authors:** Arian Amirvaresi, Arash Shahsavari, Reza Ovissipour

## Abstract

Cultivated meat production requires stable, scalable cell lines that maintain essential biological characteristics. Here, we establish and comprehensively characterize CDK4/hTERT immortalized ovine satellite cells using flow cytometry-based isolation and integrated transcriptomic-proteomic analysis. Flow cytometry identified highly purified satellite cells which were successfully immortalized and maintained stable proliferation (14-22h doubling time) through 67 passages. Multi-omics analysis revealed progressive molecular evolution with 754-1180 differentially expressed genes and 960 differentially expressed proteins, demonstrating coordinated transcriptional and post-transcriptional adaptations. Immortalization triggered species-specific extracellular matrix remodeling and inflammatory activation while preserving essential myogenic markers and apoptotic machinery. Immunofluorescence analysis demonstrated that key proteins, including PAX7, remained detectable despite transcriptional downregulation and absence of corresponding signals in mass spectrometry. Pathway analysis confirmed non-oncogenic profiles with enhanced metabolic flexibility and preserved differentiation capacity. These findings provide the first comprehensive molecular characterization of immortalized ovine satellite cells, revealing species-specific responses crucial for cultivated meat applications and establishing a framework for systematic cell line safety assessment.

## 1. Introduction

The global demand for animal protein is projected to increase by 70% by 2050, necessitating innovative approaches to sustainable meat production. Cultivated meat, produced by growing animal cells in controlled environments, offers a promising solution that could reduce environmental impact, enhance animal welfare, and improve food security while maintaining nutritional equivalence to conventional meat. However, the commercial viability of cultivated meat depends critically on establishing stable, scalable cell lines that retain essential biological characteristics while enabling cost-effective production at industrial scales^1,2^.

Satellite cells, the resident stem cells of skeletal muscle, represent the most promising cell type for cultivated meat applications due to their intrinsic myogenic programming and differentiation potential. However, primary satellite cells undergo senescence after limited population doublings, constraining their utility for large-scale production. Cellular immortalization through targeted genetic modifications offers a solution by extending proliferative lifespan while preserving cellular characteristics essential for muscle tissue formation^3,4^.

Among various immortalization strategies, the dual CDK4/hTERT approach has emerged as particularly promising for maintaining cellular authenticity. CDK4 bypasses p16-mediated senescence by promoting G1/S cell cycle transition, while hTERT prevents replicative senescence by extending telomeres. This combination avoids oncogenic transformation associated with viral immortalization methods, maintaining genomic stability and preserving differentiation capacity^3,5^.

Recent advances in cultivated meat have demonstrated the successful establishment of stable cell lines across multiple species, including bovine^3^ and porcine muscle stem cells for cultured meat production^6^, and bovine fibro-adipogenic progenitors for adipose tissue engineering^7^. Alternative approaches using porcine epiblast stem cells have generated 3D meat-like structures^8^, though satellite cells remain the physiologically relevant myogenic source. Recent breakthroughs extend beyond mammalian systems, with spontaneous immortalization of fish muscle stem cells enabling cultivated seafood production^9^. While these advances demonstrate technical feasibility, species-specific optimization remains critical, as previous sheep cell immortalization studies revealed unique molecular responses to cell cycle regulator expression^10^.

Despite these advances, the physiological and molecular effects of immortalization remain incompletely characterized in livestock species. While small-animal and human studies indicate that CDK4/hTERT immortalization preserves core muscle regulatory networks and non-oncogenic profiles^5^, emerging evidence shows that livestock cells exhibit species-specific adaptations, including extracellular matrix remodeling, metabolic reprogramming, and altered inflammatory responses^10^. Such differences underscore the necessity of species-tailored immortalization strategies and rigorous molecular characterization before these lines can be safely and effectively deployed in food systems.

Multi-omics approaches, integrating transcriptomic and proteomic analyses, provide unprecedented resolution for characterizing cellular reprogramming and identifying potential safety concerns. These comprehensive molecular profiling strategies enable systematic evaluation of pathway alterations, identification of off-target effects, and validation of preserved cellular characteristics essential for cultivated meat applications^11,12^.

Here, we establish and comprehensively characterize immortalized ovine satellite cells using flow cytometry-based isolation, CDK4/hTERT immortalization, and integrated transcriptomic-proteomic analysis. Ovine cells serve as an important model for cultivated meat development, representing a major livestock species with distinct molecular characteristics that may inform broader applications across ruminant species. Through systematic multi-omics profiling comparing primary, early-passage, and late-passage immortalized cells, we reveal progressive molecular evolution involving coordinated transcriptional and post-transcriptional adaptations.

To the best of our knowledge, this study provides the first comprehensive molecular characterization of CDK4/hTERT immortalized ovine satellite cells and establishes a framework for systematic evaluation of immortalized cell lines for cultivated meat applications, addressing critical knowledge gaps in cellular reprogramming and safety assessment that will inform future cell line development strategies.

## 2. Materials and methods

### 2.1 Primary Satellite Cell Isolation and Culture Establishment

Primary lamb satellite cells (PLSCs) were isolated using an established bovine satellite cell protocol^3^. Fresh muscle tissue was collected post-mortem from the latissimus dorsi of a 10-month-old Suffolk lamb at the Rosenthal Meat Science Center, Texas A&M University, College Station, USA. The animal was processed as part of routine operations for meat production, not specifically for this research; therefore, no additional Institutional Animal Care and Use Committee (IACUC) approval was required beyond standard institutional animal welfare protocols.

The collected sample was maintained and transported on ice in a 50 ml Falcon tube containing Dulbecco’s Modified Eagle Medium (DMEM; Thermo Fisher Scientific, cat. no. 10566024) supplemented with 1% Primocin (InvivoGen, cat. no. ant-pm-1, San Diego, CA, USA). The samples (0.7–0.8 g) were transferred into aseptic conditions in a biosafety cabinet and minced properly using sterile forceps and scissors into 10 cm organ dishes (Millipore Sigma, cat. no. CLS430165). Subsequently, the samples were transferred into 50 ml Falcon tubes (Millipore Sigma, cat. no. CLS430829) and incubated at 37°C in media containing DMEM with 0.5% collagenase II (Thermo Fisher Scientific, cat. no. 17101015) for at least 45 min on a 100 rpm shaker. Upon tissue enzymatic digestion, a 20-gauge needle was used to pass the isolated cells into the Falcon tube. The cells were resuspended in growth medium composed of DMEM supplemented with 20% fetal bovine serum (Thermo Fisher Scientific, cat. no. 26140079), 1 ng/ml human fibroblast growth factor-2 (FGF-2; Thermo Fisher Scientific, cat. no. 100-18B), and 1% Primocin. To complete isolation, the resuspended cells were passed through a 100 μm cell strainer followed by a 40 μm cell strainer (Corning, cat. no. CLS431751) and proceeded to automated cell counting processing (Thermo Fisher Scientific, Countess automated cell counter). Finally, the cells at 100,000 cells/cm² were plated onto T25 tissue culture flasks coated with 0.1% (w/v) gelatin (VWR, cat. no. 97062-618; Thermo Fisher Scientific, cat. no. 690175) in a 37°C incubator with 5% CO₂. Flasks were left for three days before the first medium change. Upon reaching 70-80% confluency, the cells were harvested using 0.25% trypsin-EDTA (Thermo Fisher Scientific, cat. no. 25200056) and passaged to a new coated flask.

### 2.2 Flow Cytometric Cell Sorting and Purification

Cell sorting was performed at the Flow Cytometry Facility, Department of Veterinary Medicine and Biomedical Sciences, Texas A&M University, College Station, USA. Frozen lamb muscle-derived cells were recovered in a 37°C water bath and washed twice with phosphate-buffered saline (PBS). Single cell suspensions were stained with DAPI to distinguish live from dead cells.

To identify and purify satellite cells, we used a flow cytometry antibody panel specifically designed to enrich muscle progenitor cells. CD31-FITC and CD45-FITC were included for negative selection, enabling the exclusion of endothelial and hematopoietic populations, respectively. For positive selection, CD56-PE-Cy7 and CD29-APC were employed as surface markers to reliably isolate satellite cells. All antibodies were incubated for 30 min at 4°C in FACS buffer. Cell sorting was performed using a Cytek Aurora 5-laser (UV 355 nm/Violet 405 nm/Blue 488 nm/Yellow-Green 561 nm/Red 640 nm) spectral flow cytometer equipped with 64 fluorescence detectors and 3 scatter channels. The instrument utilized full spectrum technology with spectral unmixing performed using Cytek SpectroFlo acquisition software to resolve overlapping fluorochrome emissions. A 100 μm nozzle tip was employed for cell sorting with optimized parameters: drop drive frequency of 24,500 Hz, amplitude of 11,087, and drop delay of 8,146 microseconds. The sort mode was set to “purity” to maximize the accuracy of sorted populations. Data acquisition was performed at a rate of less than 5,000 events per second to maintain sort accuracy and cell viability. The gating strategy systematically excluded cell debris, cell aggregates, DAPI-positive dead cells, CD31⁺ endothelial cells, and CD45⁺ hematopoietic cells, leaving only viable CD31⁻CD45⁻CD56⁺CD29⁺ satellite cells for collection. Approximately 1×10⁶ live, single, CD45⁻CD31⁻CD56⁺CD29⁺ cells were sorted into collection tubes containing Ham’s F-10 medium supplemented with 20% fetal bovine serum and maintained at 4°C throughout the procedure.

### 2.3 Cell Culture Quality Control and Contamination Testing

Mycoplasma contamination testing was carried out using MycoAlert PLUS Mycoplasma Detection Kit (Lonza, cat. no. LT07-318, Walkersville, MD, USA). Primary lamb satellite cells (PLSCs, passage 2) were cultured in triplicate to reach 70% confluency in 6-well plates. Cells were trypsinized with 1 ml of 0.25% trypsin-EDTA and neutralized with maintenance medium, then centrifuged at 300 × g for 5 min. Mycoplasma testing was performed using 100 μl of cell culture supernatant placed in a 96-well plate according to manufacturer’s instructions. Positive and negative controls were assessed using MycoAlert Assay Control Set (Lonza, cat. no. LT07-518) to validate assay performance and ensure accurate contamination detection.

### 2.4 Dual Lentiviral Immortalization Strategy

Primary lamb satellite cells were immortalized using dual lentiviral transduction with human telomerase reverse transcriptase (hTERT) and cyclin-dependent kinase 4 (hCDK4) to overcome replicative senescence while preserving myogenic characteristics. Human CDK4 was subcloned into lentiviral expression vector pLenti-CMV-MCS-EF1-puro using standard molecular cloning techniques. Lentiviral particles were produced using HEK293FT packaging cells. Briefly, 5×10⁶ HEK293FT cells were seeded in 10 cm culture dishes and co-transfected the following day with lentiviral transfer vector (pLenti-hTERT or pLenti-hCDK4. Viral supernatant collected 48 hours post-transfection, concentrated, aliquoted, and stored at -80°C. Primary lamb satellite cells (2×10⁵ cells) were transduced with hTERT lentivirus at a multiplicity of infection (MOI) of 5 and co-transduced with hCDK4 lentivirus at MOI 5 to enhance proliferative capacity while maintaining myogenic potential. The cells were selected by puromycin at the concentration of 1 ug/ml for three days and continued to passage for about 3 passages. The cells were expanded and frozen at 1 million cells per vial. Transgene integration and expression were confirmed by polymerase chain reaction (PCR) analysis using genomic DNA extracted from transduced cells cultured to 90% confluency in 6-well plates. PCR amplification was performed using hTERT forward (5’-CCAAGAACGCAGGGATGTCG-3’) and reverse (5’-AGGGCAGTCAGCGTCGTC-3’) primers yielding a 200 bp amplicon, and hCDK4 forward (5’-CTTGCCAGCCGAAACGATCA-3’) and reverse (5’-CCAGCTTGACTGTTCCACCA-3’) primers yielding a 153 bp amplicon. PCR products were analyzed by agarose gel electrophoresis, confirming successful integration of both transgenes in immortalized satellite cells compared to parental controls, and all immortalized cell lines tested negative for mycoplasma contamination using standard PCR-based detection methods. The immortalized ovine muscle satellite cells expressing both hTERT and hCDK4 transgenes were expanded and cryopreserved for subsequent experimental applications.

### 2.5 Molecular Validation of Immortalization by Quantitative PCR

To validate successful immortalization and confirm expression of transgenes in engineered satellite cells, relative gene expression analysis was performed using quantitative reverse transcription polymerase chain reaction (qPCR). Total RNA was harvested from primary and immortalized cells at 70% confluency using the RNeasy Mini Kit (Qiagen, cat. no. 74104, Hilden, Germany) according to manufacturer’s instructions. Complementary DNA (cDNA) synthesis was performed using the High-Capacity cDNA Reverse Transcription Kit (Applied Biosystems, cat. no. 4368814) with 1 μg of total RNA as template. Quantitative PCR reactions were conducted using 2 μL of cDNA template, gene-specific primers for hTERT and hCDK4 transgenes, and SYBR Green Master Mix (Bio-Rad, cat. no. 1725124) on a CFX Opus 96 Real-Time PCR System (Bio-Rad). Relative gene expression was calculated using the 2^−ΔΔCt method, normalized to β-actin as the housekeeping gene, with results expressed as fold changes relative to primary satellite cells to confirm transgene expression in immortalized cells. Each sample was analyzed in technical triplicate to ensure reproducibility.

### 2.6 Long-term Proliferative Capacity and Growth Kinetics Assessment

Following passage 3, primary and immortalized lamb cells were seeded onto T75 tissue culture flasks (Thermo Fisher Scientific, cat. no. 156800) coated with 0.1% (w/v) gelatin (VWR, cat. no. 97062-618) at a density of 7,000 cells/cm². Culture medium was replaced every 2 days until cell density reached 70-80% confluency. Upon reaching target confluency, cells were harvested using 0.25% trypsin-EDTA (Thermo Fisher Scientific, cat. no. 25200056), counted using an automated cell counter (Thermo Fisher Scientific, Countess 3 FL automated cell counter), and reseeded into new gelatin-coated T75 tissue culture flasks at the same density. To ensure maintenance of cellular integrity and validate the effectiveness of the immortalization process, comprehensive monitoring was performed at each passage to assess proliferative capacity and detect early indicators of cellular senescence. Population doublings were monitored throughout the culture period to assess proliferative capacity and detect any decline in growth rate that might indicate cellular stress or senescence onset. Cellular morphology was assessed at each passage using phase-contrast microscopy to monitor for morphological changes indicative of cellular senescence.

### 2.7 Satellite Cell Identity and Myogenic Differentiation Characterization

Functional characterization assays were performed to examine satellite cell identity and myogenic differentiation capacity in primary (PLM), early-passage immortalized (EPILM), and late-passage immortalized (LPILM) cell populations. For satellite cell identity verification, cells were examined for expression of paired box protein 7 (Pax7) that defines satellite cell stemness and regulates myogenic commitment. Cultured cells were fixed using 4% paraformaldehyde (Thermo Fisher Scientific, cat. no. J61899.AP) for 20 min at room temperature, then permeabilized with 0.5% Triton X-100 (Millipore Sigma, cat. no. T8787) for 20 min to enable antibody penetration. Following three sequential washes with DPBS, non-specific binding sites were blocked using DPBS containing 5% (v/v) goat serum (Thermo Fisher Scientific, cat. no. 16210064) for 1 hour at room temperature.

Cells were then washed with DPBS supplemented with 0.1% Tween-20 and incubated overnight at 4°C with primary antibody Pax7 (1:250 dilution; Thermo Fisher Scientific, cat. no. PA5-68506) diluted in blocking solution. After thorough washing with DPBS-Tween, cells underwent secondary blocking for 30 min before incubation with secondary antibody (anti-rabbit, 1:500 dilution; Thermo Fisher Scientific, cat. no. A-11072) for 1 hour at room temperature in darkness. Nuclear visualization was achieved through DAPI staining (1:250 dilution; Abcam, cat. no. ab104139) for 15 min. Fluorescence imaging was conducted using an Olympus CKX53 microscope to evaluate Pax7 expression patterns and nuclear localization across different cell populations.

To assess myogenic differentiation potential, satellite cell populations were subjected to directed differentiation protocols following initial expansion. Cells were cultured in standard growth conditions for 3 days before transitioning to differentiation-inducing medium consisting of equal volumes of Neurobasal (Invitrogen, cat. no. 21103049) and L15 medium (Invitrogen, cat. no. 11415064), enriched with 100 ng/mL epidermal growth factor (Sigma-Aldrich, cat. no. 100-26-500UG), 10 ng/mL insulin-like growth factor 1 (Sigma-Aldrich, cat. no. 100-34AF-100UG), and 1% antibiotic-antimycotic solution. Differentiation medium was refreshed every 48 hours throughout a 28-day differentiation period to promote myotube formation and maturation.

Post-differentiation analysis focused on detecting myogenic markers indicative of early and terminal differentiation stages. Immunofluorescence staining was performed using primary antibodies targeting myosin heavy chain (MHC; 1:20 dilution; Developmental Studies Hybridoma Bank, cat. no. MF-20) as a marker of terminally differentiated myocytes, and desmin (1:200 dilution; Abcam, cat. no. ab15200) to identify early myogenic commitment and intermediate filament organization. Primary antibodies were applied overnight at 4°C in blocking buffer, followed by incubation with appropriate fluorescent secondary antibodies (MHC detection: 1:200 dilution, Thermo Fisher Scientific, cat. no. A32723; desmin detection: 1:500 dilution, Thermo Fisher Scientific, cat. no. A11072) for 1 hour at room temperature under light protection. DAPI staining (1:250 dilution; Thermo Fisher Scientific) was employed for nuclear identification.

### 2.8 Global Transcriptomic Profiling and Analysis

Total RNA was extracted from PLM at passage 5, EPILM at passage 5, and LPILM at passage 30 using standard methods. RNA quality was assessed prior to library preparation to ensure integrity suitable for downstream analysis. For transcriptome sequencing, messenger RNA was purified from total RNA using poly-T oligo-attached magnetic beads to selectively enrich for polyadenylated transcripts. Following mRNA purification, RNA was fragmented using chemical methods, and first-strand cDNA synthesis was performed using random hexamer primers.

For strand-specific library construction, second-strand cDNA synthesis was performed using dUTP instead of dTTP to preserve directional information of the original transcripts. The sequencing library was prepared through end repair, A-tailing, adapter ligation, size selection, USER enzyme digestion for strand specificity, PCR amplification, and purification. Library quality and concentration were verified using Qubit fluorometric quantification and real-time PCR, with size distribution confirmed by Bioanalyzer analysis. Following library quality control, multiplexed libraries were pooled based on effective concentration and target data yield, then subjected to NovaSeq X Plus sequencing system by Illumina. For bioinformatics analysis, raw sequencing data in FASTQ format were processed using fastp software to generate high-quality clean reads by removing adapter sequences, poly-N containing reads, and low-quality reads. Quality metrics including Q20, Q30 scores and GC content were calculated for the filtered dataset. Reference genome and gene model annotation files were obtained from public genome databases, and HISAT2 (version 2.2.1) was used to build genome indices and align paired-end clean reads to the reference genome.

For samples requiring novel transcript discovery, mapped reads were assembled using StringTie (version 2.2.3) with a reference-based approach to identify previously unannotated splice variants and transcripts. Gene expression quantification was performed using featureCounts (version 2.0.6) to enumerate reads mapped to annotated gene features, followed by calculation of Fragments Per Kilobase of transcript per Million mapped reads (FPKM) values. FPKM normalization accounts for both sequencing depth and gene length effects, providing standardized expression measurements suitable for comparative analysis.

Differential expression analysis was conducted using DESeq2 R package (version 1.42.0) for experiments with biological replicates, implementing statistical models based on negative binomial distribution to identify significantly altered transcripts. For experiments without biological replicates, the edgeR R package (version 4.0.16) was employed with scaling normalization factors to eliminate sequencing depth differences between samples. Statistical significance thresholds were set at adjusted p-value ≤ 0.05 and |log₂(fold change)| ≥ 1, with p-value correction performed using the Benjamini-Hochberg method.

Functional enrichment analysis of differentially expressed genes was performed using clusterProfiler R package (version 4.8.1) for both Gene Ontology (GO) and Kyoto Encyclopedia of Genes and Genomes (KEGG) pathway analysis, with gene length bias correction applied.

### 2.9 Comparative Proteomic Analysis and Validation

The proteomics assay was conducted by Novogene Corporation Inc. (Sacramento, USA) using samples from PLM at passage 5 and LPILM at passage 30 to validate transcriptomic findings and assess long-term protein expression stability. For protein extraction, cell pellets were lysed in buffer containing 8 M urea (Sigma-Aldrich), 1 mM PMSF (Sigma-Aldrich), and 2 mM EDTA (Sigma-Aldrich) and sonicated for 5 min on ice using ultrasonic cell grinder (Qsonica, cat. no. Q800R3). Lysates were centrifuged (15,000 × g, 10 min, 4°C; Eppendorf, cat. no. 5430R), and protein in the resulting supernatant was quantified by BCA assay (Thermo Fisher Scientific; BSA standard, Sigma-Aldrich) on a microplate reader (Thermo Fisher Scientific, cat. no. A51119600C). For digestion, 30 µg protein was adjusted to 100 µL with 8 M urea, reduced with DL-dithiothreitol to 5 mM (37°C, 45 min; Thermo Fisher Scientific), alkylated with iodoacetamide to 11 mM (room temperature, dark, 15 min; Thermo Fisher Scientific), and diluted with 400 µL 25 mM ammonium bicarbonate (Thermo Fisher Scientific). Proteolysis was performed overnight at 37°C with 1 µL of 1 µg/µL trypsin (Promega, cat. no. V5280) and 1 µL of 1 µg/µL Lys-C (Thermo Fisher Scientific, cat. no. 90057). Peptides were acidified to pH 2–3 with 20% TFA, desalted using C18 sorbent (Millipore, Billerica, MA), and quantified with the Pierce™ Quantitative Peptide Detection Kit (Thermo Fisher Scientific).

Protein samples were analyzed using a NanoElute UHPLC system operating at a nanoliter flow rate. Mobile phase A consisted of 0.1% formic acid in water, and mobile phase B consisted of 0.1% formic acid in acetonitrile (100% ACN). Peptides (300 ng) were injected via an autosampler onto an analytical column (IonOpticks, Australia; 25 cm × 75 µm, C18, 1.6 µm particles) maintained at 50°C. Separation was performed at 300 nL/min with the following 47 min gradient: 2–35% B (0– 40 min), 35–95% B (40–40.5 min), and 95% B (40.5–47 min).

Mass spectrometric detection was performed on a timsTOF HT mass spectrometer in positive ion mode. The ddaPASEF method was first used to optimize the acquisition windows for subsequent diaPASEF runs. General MS settings included: valid gradient of 47 min, scan range 100–1700 m/z, ion mobility range (1/K₀) 0.6–1.6 Vs/cm², accumulation and release time 100 ms, ion utilization ∼100%, capillary voltage 1600 V, drying gas 3 L/min at 180°C. For ddaPASEF, 10 PASEF MS/MS frames were acquired per cycle (1.17 s), with a target intensity of 20,000, threshold 2500, charge states 0–5, and collision energy ramped from 59 eV at 1/K₀ = 1.6 Vs/cm² to 20 eV at 1/K₀ = 0.6 Vs/cm². Quadrupole isolation widths were 2 Th for m/z < 700 and 3 Th for m/z > 800. For diaPASEF acquisition, settings included a mass range of 400–1200 m/z, mobility range 0.6–1.6 Vs/cm², 25 Da isolation windows with 0.1 Da overlap, 24 mass steps per cycle, and 2 mobility windows, totaling 48 acquisition windows. The average cycle time was 1.17 s.

Mass spectrometry data were processed using DIA-NN (v1.8.1) software for database searching against ovine protein databases (uniprotkb_proteome_UP000002356_sheep_2024_09_25.fasta, 23,108 sequences) with library-free method and deep learning-based spectral library prediction. Match-between-runs was enabled to improve protein quantification, with false discovery rates filtered at 1% for both precursor and protein level identification. Protein quantification was performed using the MaxLFQ algorithm with median-based normalization, applying centroid transformation for relative quantitative values. Quality control assessment included evaluation of peptide length distribution, missed cleavage sites, and inter-sample correlation analysis using Pearson correlation coefficients. Differential expression analysis was conducted using t-tests for two-group comparisons with significance criteria of fold change ≥ 1.5 or ≤ 0.667 combined with P-value ≤ 0.05. Multiple testing correction was applied using the Benjamini-Hochberg method to control false discovery rate. Principal component analysis and hierarchical clustering with z-score normalization were performed to assess sample relationships and identify expression patterns. Functional enrichment analysis was conducted using Gene Ontology and KEGG pathway databases.

#### Statistical analysis

All statistical analyses were performed using GraphPad Prism (version 10.0; GraphPad Software, San Diego, CA, USA). For comparison between two groups, unpaired two-tailed Student’s t-tests were used.

## 3. Results

### 3.1 Cell Sorting and Establishment of Immortalized Ovine Satellite Cells

#### 3.1.1 Flow cytometric characterization and satellite cell isolation

Flow cytometry analysis demonstrated successful isolation and enrichment of ovine satellite cells from primary muscle tissue using a comprehensive sequential gating strategy (Figure 1a-e). Initial gating excluded cellular debris based on forward and side scatter characteristics (FSC-A vs. SSC-A), capturing 95.05% of total events as intact cells for subsequent analysis (Figure 1a). Doublet discrimination using scatter height versus area plots (SSC-H vs. SSC-A) identified 99.24% of cellular events as singlets (Figure 1b), ensuring accurate single-cell quantification. Viability assessment using DAPI exclusion revealed 98.32% live cells within the singlet population (Figure 1c), indicating excellent tissue preservation and gentle processing conditions.

**Figure 1.**
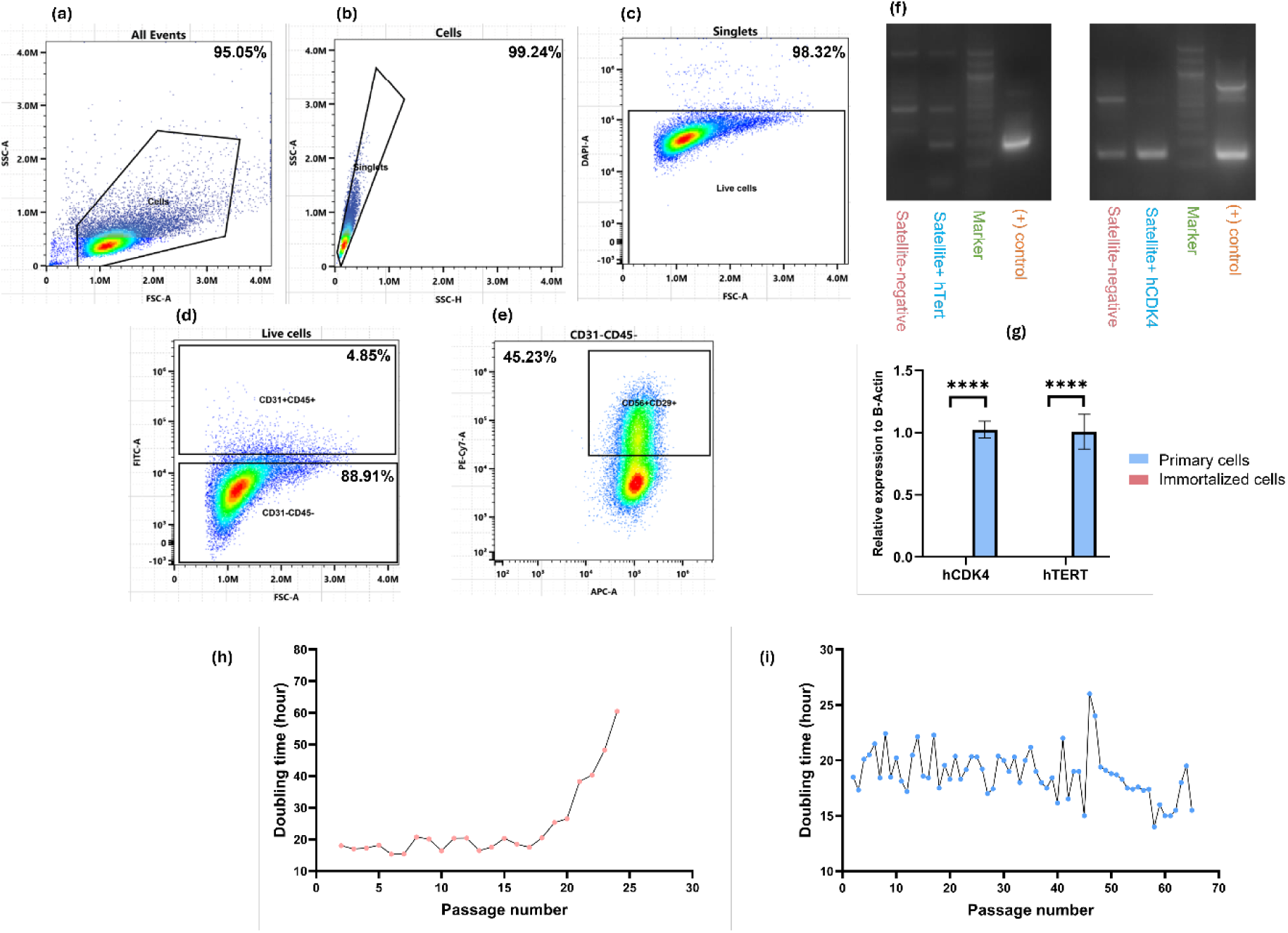
Flow cytometric isolation and functional validation of immortalized ovine satellite cells. (a-e) Sequential gating strategy for satellite cell purification from primary ovine muscle tissue. (a) Forward scatter (FSC-A) vs. side scatter (SSC-A) plot showing initial cell population with debris exclusion gate capturing 95.05% of cellular events. (b) Doublet discrimination using SSC-H vs. SSC-A, identifying 99.24% singlet cells. (c) Live/dead cell discrimination using DAPI staining, showing 98.32% cell viability in the singlet population. (d) Negative selection gate for CD31⁻CD45⁻ cells (88.91% of live cells), excluding endothelial and hematopoietic cell contamination. (e) Positive selection for satellite cell markers CD56⁺CD29⁺ within the CD31⁻CD45⁻ population, identifying 45.23% double-positive cells. (f) Agarose gel electrophoresis confirming successful genomic integration of hTERT and hCDK4 transgenes. Left panel shows satellite-negative control, satellite+hTERT, marker, and positive control lanes. Right panel shows satellite-negative control, satellite+hCDK4, marker, and positive control lanes, demonstrating specific amplification of transgenes in immortalized cells. (g) Quantitative PCR validation of transgene expressions showing significant upregulation of hCDK4 and hTERT relative to β-actin in immortalized cells compared to primary cells (n=3 biological replicates, ****p<0.0001, unpaired t-test). (h-i) Long-term growth kinetics analysis comparing population doubling times. (h) Primary lamb muscle cells (PLM) showed stable doubling times (16-22 hours) until passage 19, followed by senescence-associated growth decline reaching ∼60 hours by passage 26. (I) Immortalized lamb muscle cells (ILM) maintained consistent doubling times (14-22 hours) across 67+ passages with transient fluctuations but no progressive senescence.

The critical purification steps involved sequential negative and positive selection strategies to enrich for myogenic progenitors. Based on CD31 and CD45 expression, two distinct populations were identified (Figure 1d). The majority of live cells (88.91%) exhibited a CD31⁻CD45⁻ phenotype, representing non-endothelial and non-hematopoietic muscle-resident cells. A small subset (4.85%) was positive for CD31⁺CD45⁺, indicating contaminating endothelial and hematopoietic cells co-isolated during tissue digestion, which were effectively removed through negative selection. This depletion strategy is well-established across species, with studies showing CD31/CD45 negative selection effectively removes vascular and immune cell contaminants while preserving satellite cell viability and function ^13–16^.

Further analysis of the CD31⁻CD45⁻ population for satellite cell-specific markers revealed that 45.23% of cells expressed both CD56⁺CD29⁺ (Figure 1e), representing a substantial enrichment of satellite cell populations. CD56 (NCAM) and CD29 (integrin β1) are established satellite cell surface markers that have been validated in bovine and porcine skeletal muscle systems^14,16^. The observed 45.23% CD56⁺CD29⁺ expression within the CD31⁻CD45⁻ population is consistent with published studies in other livestock species. Similar findings have been reported in porcine muscle stem cells, where CD29 was exclusively expressed in muscle stem cells while CD56 single-positive cells represented approximately 25% of the population, with CD56⁺CD29⁺ cells comprising the majority of myogenic cells after CD29-based purification^17^.

#### 3.1.2 Successful immortalization through dual genetic modification

Ovine muscle satellite cells were successfully immortalized through dual genetic modification using separate lentiviral vectors encoding human TERT and CDK4 (Figure 1f-g). Our lentiviral CDK4/hTERT approach builds upon previous sheep cell immortalization strategies using cell cycle regulators^10^ but provides comprehensive multi-omics characterization specifically for satellite cells. This immortalization strategy was selected based on established mechanisms where hTERT extends telomeres to prevent replicative senescence, while hCDK4 promotes cell cycle progression through G1/S transition by bypassing p16-mediated growth arrest. Studies by Thorley et al. demonstrated that bypassing both pathways is required for complete immortalization, with neither hTERT nor CDK4 alone being sufficient^5^.

Gel electrophoresis analysis confirmed successful genomic integration of both transgenes in immortalized cells (Figure 1f). The left panel demonstrates specific amplification of hTERT in satellite+hTERT samples compared to satellite-negative controls, with no detectable band in the negative control lane, confirming transgene-specific expression. The right panel shows hCDK4 amplification results, where the satellite-negative control exhibits faint banding, likely representing basal endogenous CDK4 expression in ovine cells. However, the immortalized satellite+hCDK4 sample displays a substantially stronger and more intense band at the expected size (∼153 bp), indicating successful overexpression of the human CDK4 transgene. Both panels include molecular weight markers and positive controls validating the PCR amplification specificity. The hTERT amplification (∼200 bp) showed complete specificity with no background in control samples, while the hCDK4 results demonstrate clear differential expression between baseline endogenous levels and transgene-driven overexpression.

Quantitative PCR analysis confirmed significant upregulation of both transgenes in immortalized cells, with hCDK4 and hTERT showing >100-fold increased expression relative to β-actin compared to primary cell controls (*p<0.001, Figure 1g). The qPCR data shows that immortalized cells exhibited relative expression values of approximately 1.0 for both hCDK4 and hTERT when normalized to β-actin, while primary cells showed no detectable expression above baseline. All cell cultures tested negative for mycoplasma contamination, and immortalized cells demonstrated successful cryopreservation with >90% post-thaw viability, indicating establishment of a stable, bankable cell line resource.

#### 3.1.3 Enhanced proliferative capacity and extended lifespan

The proliferative capacity evaluation revealed dramatic differences between primary and immortalized ovine satellite cells (Figure 1h-i). Primary ovine satellite cells maintained relatively stable doubling times between 16-22 hours up to passage 19, consistent with normal satellite cell behavior in early passages (Figure 1h). However, beyond passage 19, primary cells showed significant increases in doubling time, reaching approximately 40 hours by passage 21 and accelerating to ∼60 hours by passage 26 due to cellular senescence resulting from telomere shortening and cell cycle arrest.

In contrast, immortalized ovine satellite cells maintained significantly more stable proliferative properties over an extended passage range (Figure 1i, extending to passage 67 and ongoing), with doubling times consistently fluctuating between 15-22 hours with no significant difference between early (P5-15) and late passages (P50-67). Throughout the 67+ passages monitored, immortalized cells showed minor fluctuations in doubling time. Notably, doubling times at late passages (P60-67) remained between 15-20 hours, suggesting maintained proliferative vigor without signs of growth crisis or replicative exhaustion. This proliferative stability represents a substantial improvement over primary cells, which typically senesce after 15-25 population doublings in our culture conditions.

Our results align closely with previous successful livestock satellite cell immortalization studies. Stout et al. reported successful bovine satellite cell immortalization using hTERT and CDK4, achieving 15-hour doubling times across more than 120 passages. Despite technical differences in gene delivery methods , our study employed lentiviral vectors (more accessible and widely used in primary cell engineering) versus their Sleeping Beauty transposon system (offering integration specificity and genomic stability) , both approaches effectively delivered transgenes and resulted in stable, transgene-positive populations suitable for long-term culture and cryopreservation with comparable growth kinetics and cellular characteristics ^3^. However, as demonstrated in our subsequent analyses (Sections 3.2-3.3), species-specific molecular evolution occurs in lamb cells that was not observed in human studies, emphasizing the importance of comprehensive molecular characterization even with validated immortalization approaches.

### 3.2 Progressive molecular evolution during CDK4/hTERT immortalization reveals species-specific adaptations

To investigate the molecular consequences of CDK4/hTERT immortalization in lamb muscle cells, we performed comprehensive RNA sequencing analysis comparing primary lamb muscle (PLM; Passage 5), early-passage immortalized lamb muscle (EPILM; passage 5), and late-passage immortalized lamb muscle (LPILM; passage 30) cells.

#### 3.2.1 Global transcriptomic profiling reveals distinct molecular states

Multi-dimensional transcriptomic analysis demonstrated excellent data quality and clear molecular separation between cell populations. Hierarchical clustering revealed distinct expression patterns, with samples clustering by cell type (PLM, EPILM, LPILM) and showing characteristic gene expression signatures for each population (Figure 2a). FPKM distribution analysis showed consistent bimodal patterns across all samples, with characteristic peaks at low expression (0-1 FPKM) representing silent/lowly expressed genes and broader distributions at higher values reflecting active transcription (Figure 2b), confirming robust RNA-seq data quality. Principal component analysis revealed distinct transcriptomic signatures, with PC1 (44.9% variance) and PC2 (26.6% variance) effectively separating the three muscle cell populations into well-defined clusters (Figure 2c). The spatial arrangement revealed that early immortalization induced substantial molecular changes separating EPILM from PLM, while extended passage drove LPILM even further along PC1, indicating progressive, directional molecular evolution.

**Figure 2.**
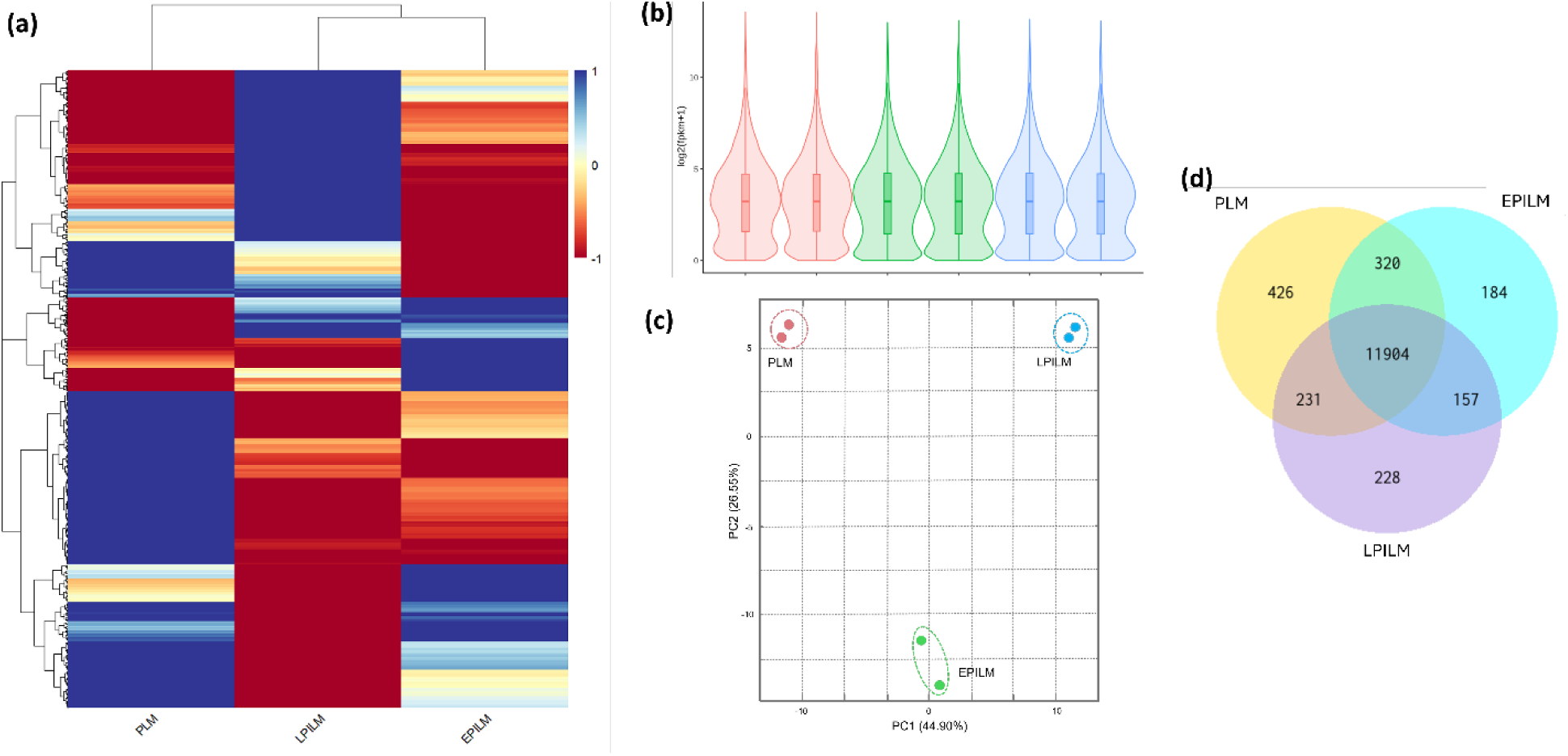
Transcriptomic characterization of muscle cell populations. (a) Hierarchical clustering heatmap showing gene expression patterns across PLM (primary lamb muscle), EPILM (early passage immortalized lamb muscle), and LPILM (late passage immortalized lamb muscle) samples. (b) Violin plots displaying FPKM (Fragments Per Kilobase of transcript per Million mapped reads) distribution across all samples, showing characteristic bimodal patterns with peaks at low expression levels and broader distributions for highly expressed genes. (c) PCA plot showing clear separation of the three muscle cell populations based on their transcriptomic profiles. Each dot represents an individual sample, with colors indicating cell type. (d) Venn diagram illustrating the overlap of differentially expressed genes between PLM, LPILM, and EPILM conditions with 11904 genes commonly expressed across all comparisons and unique gene sets for each condition.

Differential gene expression analysis identified 11,904 commonly expressed genes across all conditions, with condition-specific gene sets comprising 426 PLM-specific, 184 EPILM-specific, and 228 LPILM-specific genes (Figure 2d). The overlap patterns indicated that while a core transcriptional program is maintained, each immortalization state exhibits unique molecular features, with progressive loss of condition-specific genes from primary (426) through early (184) to late (228) passage, suggesting convergence toward a stable immortalized transcriptional state.

#### 3.2.2 Stage-specific molecular reprogramming intensifies with passage progression

Comprehensive differential expression analysis revealed progressive molecular evolution with distinct characteristics at each immortalization stage. Early immortalization (EPILM vs PLM) involved 754 differentially expressed genes (DEGs; |log_2_FC| > 1, padj < 0.05) with pronounced transcriptional suppression (67.2% downregulated), late immortalization (LPILM vs PLM) encompassed 1,180 DEGs maintaining suppressive bias (63.5% downregulated), while passage progression (LPILM vs EPILM) showed 818 DEGs with more balanced regulation (53.4% downregulated), indicating adaptive responses during extended culture (Figure 3a, d, g).

**Figure 3.**
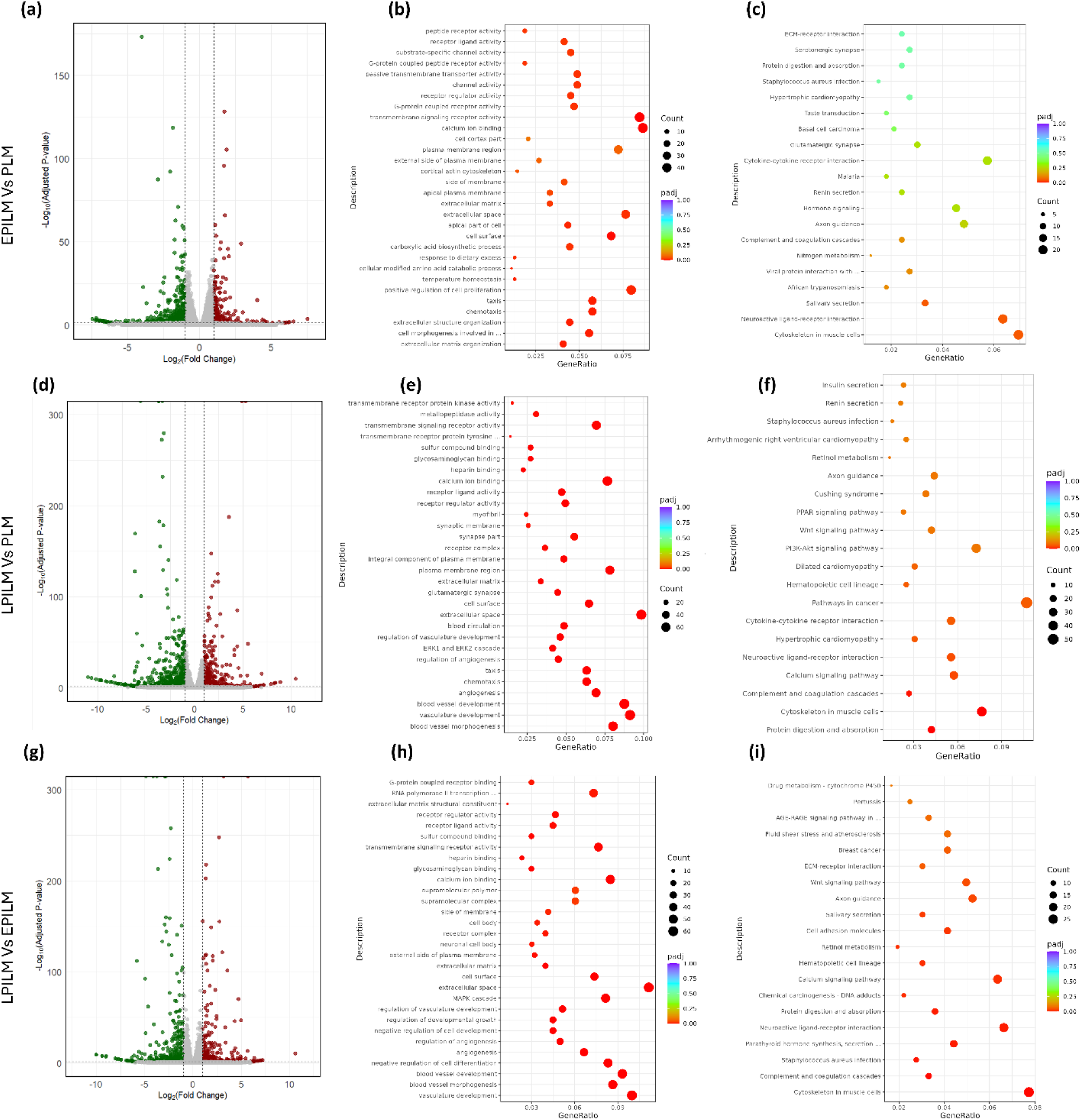
Comprehensive differential expression and pathway enrichment analysis reveals progressive molecular evolution across immortalization stages. (a, d, g) Volcano plots showing differentially expressed genes for EPILM vs PLM (a, 754 DEGs), LPILM vs PLM (d, 1180 DEGs), and LPILM vs EPILM (g, 818 DEGs). Green dots indicate downregulated genes, red dots indicate upregulated genes (|log2FC| > 1, padj < 0.05). Progression shows increasing fold-change magnitudes and more extreme alterations. (b, e, h) Gene Ontology enrichment bubble plots showing significantly altered categories. EPILM vs PLM (b) shows 43 enriched categories with ECM and developmental processes; LPILM vs PLM (e) demonstrates 511 categories with prominent vascular development; LPILM vs EPILM (h) reveals 621 categories with stress response emergence. Bubble size indicates gene count, color indicates significance. (c, f, i) KEGG pathway enrichment bubble plots showing nominally significant pathways for EPILM vs PLM (c, 0 pathways achieving padj < 0.05) and significantly altered pathways for LPILM vs PLM (f, 6 pathways, padj < 0.05) and LPILM vs EPILM (i, 14 pathways, padj < 0.05). Statistical significance emerges only with extended passage, demonstrating progressive coordination from scattered adaptations to network-level reorganization.

##### 3.2.2.1 Early immortalization: Targeted molecular adaptation

Volcano plot analysis of EPILM vs PLM revealed moderate fold changes (predominantly |log_2_FC| 1-4) with 507 downregulated and 247 upregulated genes (Figure 3a). Gene Ontology enrichment analysis demonstrated statistically robust functional reorganization across 43 significantly enriched categories, with extracellular matrix organization (padj = 0.007, 22 genes with 15 downregulated), cell morphogenesis involved in neuron differentiation (padj = 0.007, 30 genes predominantly downregulated), and extracellular structure organization (padj = 0.007, 24 genes) as the top biological processes (Figure 3b). Molecular function analysis revealed calcium ion binding as most significantly enriched (46 genes, 32 downregulated/14 upregulated), indicating altered calcium homeostasis critical for muscle contraction and cellular signaling^18^. Cellular component changes focused on extracellular space and cell surface remodeling (Figure 4a), reflecting altered cell-matrix interactions characteristic of immortalized cells^19^. The predominant downregulation across functional categories (visible in Figure 4a green bars exceeding red bars) confirms that early immortalization primarily involves suppression of primary cell characteristics rather than activation of new programs.

**Figure 4.**
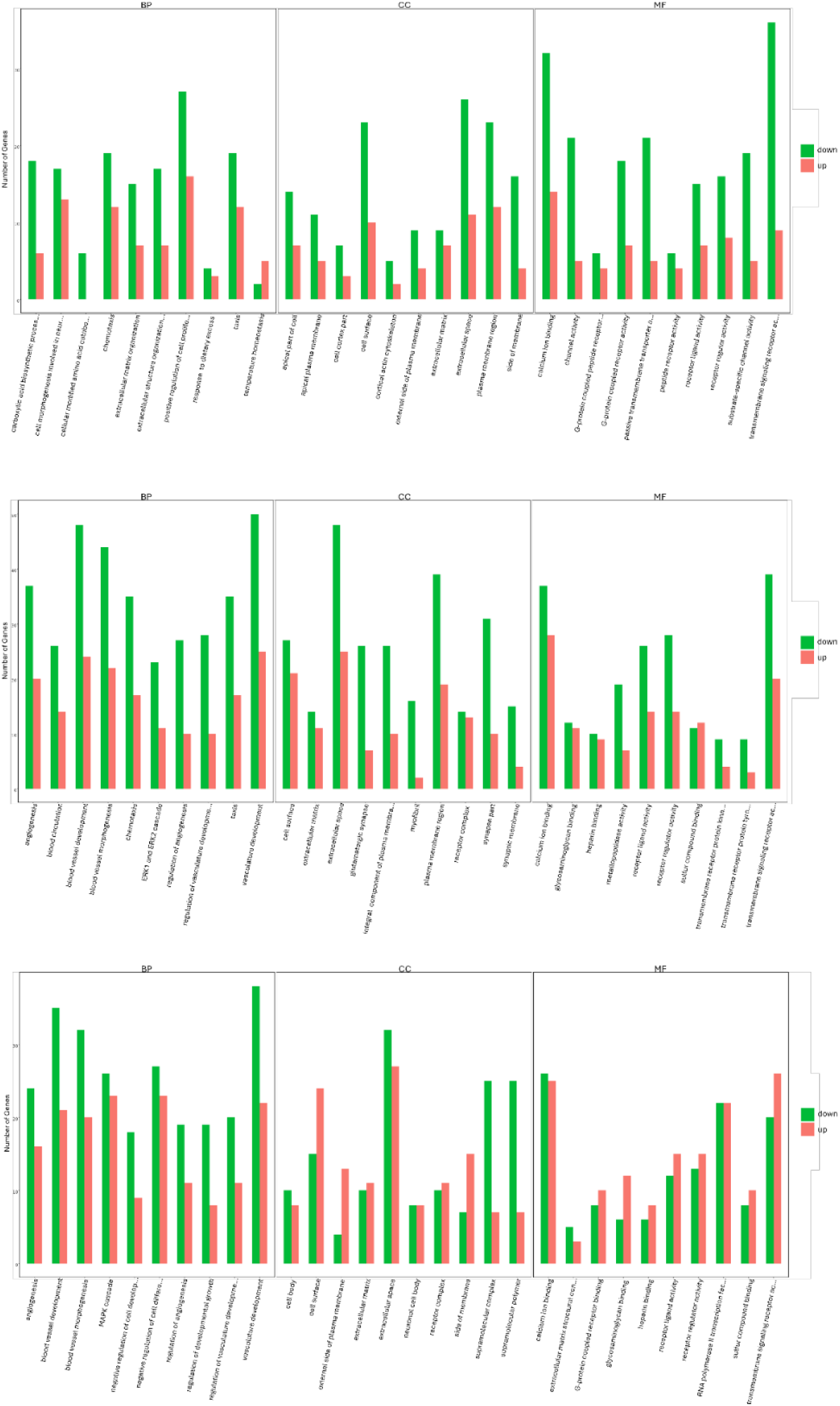
Gene Ontology enrichment analysis quantifies progressive functional divergence. Bar charts showing up-regulated (red) and down-regulated (green) gene counts within significantly enriched GO terms for (a) EPILM vs PLM , (b) LPILM vs PLM, and (c) LPILM vs EPILM, organized by biological processes (BP), cellular components (CC), and molecular functions (MF). Bar height represents gene count per category. Progressive patterns include: (a) 43 enriched categories with predominant downregulation (green > red); (b) 511 categories (>10-fold increase) with larger gene counts per category and expanded vascular/metabolic involvement; (c) 621 categories (>14-fold increase from early) with most balanced directional regulation and emergence of stress response categories. The escalating number of enriched terms (43 → 511 → 621) quantitatively validates progressive molecular evolution from scattered adaptations to coordinated network reorganization.

While KEGG pathway analysis revealed nominally significant enrichment of ECM-receptor interaction (15 genes, p = 0.0004), focal adhesion (18 genes, p = 0.001), and PI3K-Akt signaling (22 genes, p = 0.003), none achieved statistical significance after multiple testing correction (all padj > 0.05), with cytoskeleton in muscle cells approaching marginal significance (padj = 0.051) (Figure 3c).

##### 3.2.2.2 Late immortalization: Intensified molecular disruption with expanded functional involvement

The LPILM vs PLM comparison demonstrated substantially greater molecular disruption, with 749 downregulated and 431 upregulated genes exhibiting larger fold changes, including multiple genes with |log_2_FC| > 8 (Figure 3d). GO analysis revealed dramatically expanded functional category involvement, 511 significantly enriched GO terms compared to 43 in early immortalization (>10-fold increase), with enhanced metabolic process alterations (5.79% enrichment in calcium ion binding), intensified cytoskeletal organization changes (cytoskeleton in muscle cells: 33 genes, padj = 8.9×10⁻⁶), and broader regulatory pathway modifications including negative regulation of cell population proliferation (2.81% enrichment) (Figure 3e).

Notably, vascular development processes emerged as dominant signatures: blood vessel morphogenesis, angiogenesis, and vasculature development each showed 20-40 genes per category, substantially larger than early-stage categories, with complex directional patterns (Figure 4b). The bar chart distribution in Figure 4b reveals both intensification (taller bars) and diversification (more categories) compared to Figure 4a, quantitatively demonstrating escalating molecular complexity. KEGG analysis identified 6 pathways achieving significance after multiple testing correction, including cytoskeleton organization, metabolic pathways, and signaling cascades (Figure 3f).

##### 3.2.2.3 Passage progression: Most extensive functional reorganization and stress response emergence

Comparison of LPILM vs EPILM revealed 818 DEGs representing changes specifically attributable to extended passage (Figure 3g). GO enrichment identified the most extensive functional remodeling of any comparison, 621 significantly enriched GO terms, representing a further 20% increase over the late vs primary comparison and >14-fold increase over early immortalization, with novel processes not prominent in earlier comparisons, particularly stress response mechanisms (cellular response to lipopolysaccharide, response to chemical stimuli, response to glucose), drug metabolism pathways, and inflammatory signaling (Figure 3h, Figure 4c)^20,21^. The directional analysis in Figure 4c reveals a striking shift toward more balanced up/down regulation compared to the suppression-dominant patterns in Figures 4a and 4b, indicating that passage progression involves active transcriptional responses rather than continued suppression. The emergence of cellular component categories related to organelle stress and molecular function categories related to xenobiotic metabolism suggests cumulative cellular stress and compensatory adaptation. Remarkably, KEGG analysis revealed 14 significant pathways (Figure 3i), more than doubling the pathway involvement seen in late vs primary comparison, with prominent enrichment of inflammatory signaling (cytokine-cytokine receptor interaction: 25 genes, padj = 0.001), chemical carcinogenesis-ROS (18 genes, padj = 0.003), and drug metabolism pathways (12 genes, padj = 0.009), indicating cumulative cellular stress and compensatory adaptation during prolonged culture. The escalating functional involvement quantified by GO enrichment—from 43 terms (EPILM vs PLM) to 511 terms (LPILM vs PLM) to 621 terms (LPILM vs EPILM)—combined with the progressive emergence of statistically significant KEGG pathways (0 → 6 → 14 pathways) demonstrates that immortalization triggers initially scattered molecular changes that progressively coalesce into coordinated cellular reprogramming intensifying with passage number. This progression pattern has not been reported in human immortalized muscle cells and suggests species-specific vulnerabilities in livestock satellite cells^5,22^.

#### 3.2.3 Cell cycle regulation reveals compensatory suppression beyond targeted immortalization

The CDK4/hTERT immortalization strategy specifically targets cell cycle regulation by bypassing p16-mediated senescence and preventing telomere shortening (hTERT). CDK4 associates with D-type cyclins to drive G1/S progression, while hTERT maintains telomere length, theoretically enabling indefinite proliferation without broadly disrupting cell cycle control^4^. Our transcriptomic analysis revealed more complex downstream consequences.

CDKN1A (p21), a master cell cycle inhibitor that binds cyclin-CDK complexes to block cell cycle progression and induce differentiation^23,24^, demonstrated progressive, stage-specific suppression: undetectable in EPILM vs PLM, significant downregulation in LPILM vs PLM (log_2_FC = -1.03, padj = 9.96×10⁻¹¹), and further decline during passage progression (log_2_FC = -1.43, LPILM vs EPILM, padj = 2.66×10⁻³¹). This pattern suggests that while CDK4 initially bypasses p16 without affecting p21, prolonged culture triggers compensatory suppression of additional cell cycle inhibitors, potentially through epigenetic mechanisms or secondary pathway activation.

CDKN1C (p27), another CDK inhibitor critical for myoblast cell cycle exit, showed contrasting kinetics with early suppression (log2FC = -1.67, EPILM vs PLM) partially maintained in late passage (log2FC = -1.01, LPILM vs PLM), suggesting distinct regulatory mechanisms. CCND1 (Cyclin D1) exhibited unexpected late-stage downregulation (log_2_FC = -1.09, LPILM vs PLM; log_2_FC = -1.47, LPILM vs EPILM) despite no detectable early changes. This finding contradicts previous CDK4 immortalization studies reporting enhanced cyclin D1 expression and suggests negative feedback regulation of endogenous cyclin D1 in the presence of constitutive exogenous CDK4 activity, a species-specific compensatory mechanism not observed in human cells.

Critically, TP53 and RB1 showed no significant transcriptional changes across all comparisons, confirming that immortalization bypasses rather than inactivates these tumor suppressors, preserving genomic stability checkpoints essential for food safety applications^4,25^. This mechanistic distinction from oncogenic immortalization approaches (SV40, E6/E7) was validated by pathway analysis showing cancer pathways were predominantly downregulated (24 downregulated vs 15 upregulated genes), supporting a non-oncogenic cellular state.

#### 3.2.4 Extensive ECM remodeling represents unexpected species-specific response

Despite CDK4/hTERT’s targeted mechanism, elastin (ELN) emerged as the most dramatically suppressed gene in early immortalization (log_2_FC = -4.03, padj = 1×10⁻¹⁷³), with sustained suppression in late passage (log_2_FC = -3.26, LPILM vs PLM). Elastin provides tissue elasticity through crosslinked tropoelastin multimers, and its severe downregulation was accompanied by coordinated collagen suppression including COL1A1 (early: log_2_FC = -1.00; late: undetected), COL16A1 (early: log_2_FC = -1.88; late: log_2_FC = -3.75), COL11A2 (progressive suppression), establishing comprehensive ECM dismantling.

GO enrichment confirmed extracellular structure organization as the most significantly enriched process in early immortalization (19 of 22 ECM-related proteins downregulated, padj = 0.007), while KEGG analysis showed ECM-receptor interaction (15 genes, p = 0.0004) and focal adhesion (18 genes, p = 0.001) as nominally significant but not achieving significance after multiple testing correction (Figure 3c). Despite lack of statistical significance at the pathway level, the dramatic individual gene changes and robust GO enrichment (represented prominently within the 43 significant GO terms) confirm biologically meaningful ECM remodeling during early immortalization. Late-passage GO analysis showed intensified ECM disruption with 4.95% enrichment in extracellular space categories among the 511 significant GO terms (Figure 3e), and KOG analysis revealed 22 significant hits in extracellular structures (representing 28% of all significant KOG categories in proteomic analysis), validating transcriptomic findings at the protein level.

This extensive ECM remodeling contrasts sharply with human CDK4/hTERT immortalized muscle cells, which maintain stable ECM expression patterns through >100 passages. The species-specific response may reflect differences in: (1) baseline ECM organization between human and ovine muscle, (2) differential p21-CDK4-ECM regulatory circuits, or (3) species-specific epigenetic responses to indefinite proliferation. Recent studies showing p21 regulates ECM components in a CDK4-dependent manner support mechanistic linkage between cell cycle manipulation and structural protein expression. The progressive ECM suppression may also reflect selective pressure in culture, as ECM synthesis is metabolically costly and cells reducing ECM production may have proliferative advantages in 2D culture lacking mechanical constraints present in vivo^5,26–29^.

This finding has critical implications for cultivated meat applications, while CDK4/hTERT immortalization preserves myogenic programming, species-specific ECM alterations may affect tissue mechanical properties, scaffold interactions, and ultimately meat texture—requiring species-optimized immortalization protocols or ECM supplementation strategies.

#### 3.2.5 Progressive myogenic identity erosion despite preserved contractile machinery

Core myogenic regulatory factors exhibited dramatic, stage-dependent suppression. MYOD1, the master myogenic regulator capable of converting fibroblasts to muscle, showed moderate early suppression (log_2_FC = -3.59, EPILM vs PLM) progressing to severe late-stage downregulation (log_2_FC = -10.24, LPILM vs PLM, representing >1000-fold reduction), with additional passage-dependent decline (log_2_FC = -6.64, LPILM vs EPILM). PAX7, the satellite cell specification factor maintaining stem cell state while preventing terminal differentiation, followed similar kinetics, mild early suppression (log_2_FC = -1.83) to dramatic late downregulation (log_2_FC = -9.73, LPILM vs PLM), with continued decline during passage progression^30^ (log_2_FC = -7.89, LPILM vs EPILM). Similarly, MYOD1 mRNA was significantly downregulated in our transcriptomic analysis.

GO analysis confirmed systematic downregulation of myogenic transcription factors, with helix-loop-helix DNA-binding domain superfamilies and homeobox domain superfamilies showing significant enrichment in downregulated proteins (InterPro domain analysis), representing architectural-level collapse of early myogenic commitment programs.

#### 3.2.6 Inflammatory activation reveals cumulative cellular stress

While early immortalization showed controlled inflammatory activation, with IL6 (interleukin 6) moderately upregulated (log2FC = 1.90, EPILM vs PLM), late passage exhibited extraordinary inflammatory escalation with S100A8 upregulation >1000-fold (log_2_FC = 10.49, LPILM vs PLM; log_2_FC = 10.59, LPILM vs EPILM), representing the most dramatically upregulated gene across all comparisons. S100A8 forms the S100A8/A9 heterodimer (calprotectin), functioning as a damage-associated molecular pattern (DAMP) that activates TLR4 to induce pro-inflammatory cytokines (TNF-α, IL-6) and represents cellular stress signaling^31,32^.

GO enrichment in LPILM vs EPILM revealed cellular response to lipopolysaccharide and inflammatory response among the 621 significantly enriched GO terms (Figure 3h, Figure 4c), while KEGG analysis identified cytokine-cytokine receptor interaction (25 genes, padj = 0.001), chemokine signaling, and inflammatory pathways significantly enriched (Figure 3i).

This progressive inflammatory activation, despite CDK4/hTERT’s targeted mechanism avoiding oncogene-induced stress, indicates that sustained proliferation beyond normal senescence checkpoints triggers compensatory inflammatory programs, potentially through accumulating DNA damage, telomere dysfunction-independent pathways, or epigenetic drift. The inflammatory signature resembles senescence-associated secretory phenotype (SASP) components, suggesting that immortalization delays but does not eliminate cellular stress responses, with implications for long-term culture stability and potential immunogenicity in cultivated meat applications.

#### 3.2.7 Metabolic reprogramming supports enhanced proliferative demands

IGFBP2, facilitates proliferation through IGF-dependent and independent mechanisms, showed late-stage-specific severe downregulation (log_2_FC = -3.41, LPILM vs PLM; log_2_FC = -3.75, LPILM vs EPILM) with no detectable early changes.

GO analysis revealed calcium ion binding (5.79% enrichment) and carbonate dehydratase activity among top molecular functions within the 511 significantly enriched GO terms (Figure 3e, Figure 4b), correlating with proteomic upregulation of carbonic anhydrase isoforms CA2 and CA8 (>10-fold), which support pH homeostasis under intense glycolysis. While early-stage KEGG analysis did not achieve statistical significance for metabolic pathways (padj > 0.05), late-stage analysis showed progressive metabolic pathway involvement: 15 metabolically-related genes in LPILM vs PLM, 22 in LPILM vs EPILM, including glycolysis/gluconeogenesis with coordinated upregulation of phosphoglucomutase, aldose 1-epimerase, fructose-bisphosphate aldolase, and enolase, while 6-phosphofructokinase was downregulated—suggesting metabolic control with enhanced downstream flexibility and biosynthetic capacity.

### 3.3 Proteomic Validation Confirms Transcriptomic Predictions and Reveals Post-Transcriptional Complexity

To validate transcriptomic findings at the functional protein level and identify post-transcriptional regulatory mechanisms masked by RNA-seq analysis, we performed comprehensive mass spectrometry-based proteomic analysis comparing LPILM with PLM cells (n=3 biological replicates per condition). This dual-omics approach provides crucial validation of mRNA-level predictions while revealing protein stability, translation efficiency, and post-translational regulatory mechanisms that may not be apparent from RNA sequencing alone.

#### 3.3.1 Global proteomic profiling confirms extensive molecular reprogramming

Mass spectrometry-based proteomic analysis revealed substantial molecular reprogramming, with 960 differentially expressed proteins (DEPs) identified from 8094 detected proteins. Hierarchical clustering demonstrated robust separation of sample groups, with LPILM (LPILM1-3) and PLM (PLM1-3) forming distinct clusters in both the dendrogram and heatmap, confirming that protein-level changes mirror the transcriptomic molecular divergence observed in late-passage immortalization (Figure 5a). The clear group separation validates that transcriptomic changes translate to functional proteomic alterations.

**Figure 5.**
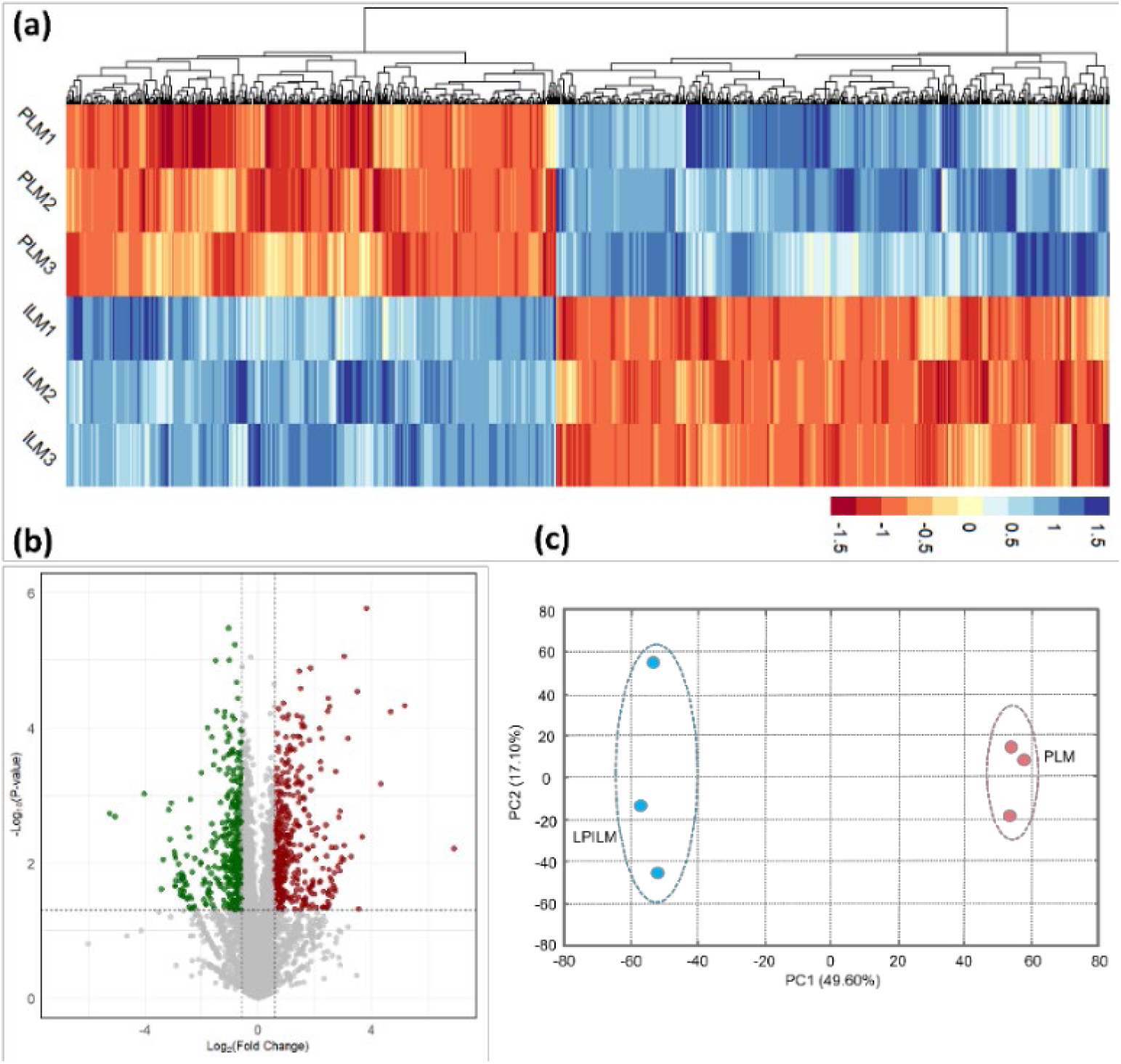
Multi-dimensional proteomic profiling validates extensive molecular reprogramming during CDK4/hTERT immortalization. (a) Hierarchical clustering heatmap showing 960 differentially expressed proteins effectively separating LPILM (late passage immortalized lamb muscle) and PLM (primary lamb muscle) samples with distinct proteomic signatures, confirming robust group separation and validating transcriptomic predictions of molecular divergence. (b) Volcano plot displaying symmetric distribution of DEPs with 451 upregulated (red) and 511 downregulated (green) proteins, demonstrating balanced proteomic reorganization that contrasts with asymmetric transcriptomic suppression and suggests compensatory post-transcriptional mechanisms. (c) Principal component analysis achieving proteomic discrimination with PC1 (49.6%) and PC2 (17.1%) separating populations into non-overlapping clusters, with higher PC1 variance than transcriptomic PCA suggesting enhanced coordination at protein level through selective translation and post-translational regulation.

PCA shows striking proteomic discrimination, with PC1 (49.6% variance) and PC2 (17.1% variance) effectively separating LPILM from PLM samples into non-overlapping clusters (Figure 5c), demonstrating that protein-level molecular reorganization is as profound as transcriptomic changes.

Furthermore, the volcano plot revealed symmetric distribution of DEPs, with 451 significantly upregulated proteins and 511 downregulated proteins in LPILM cells compared to PLM (Figure 5b).

#### 3.3.2 Transcriptome-proteome correlation reveals high concordance with strategic discordances

Our observation revealed excellent overall concordance, showing directionally concordant alterations with their corresponding mRNAs where comparable data existed. This high concordance rate validates that transcriptomic changes generally translate to functional protein-level alterations, confirming the biological relevance of RNA-seq findings while also identifying specific targets of post-transcriptional control that add regulatory complexity beyond transcriptional mechanisms.

Concordant suppression validates muscle identity collapse. MYOD1 and MYOG demonstrated concordant downregulation at both mRNA and protein levels, providing definitive validation that the dramatic transcriptomic suppression of master myogenic regulators translates to functional protein depletion. This protein-level confirmation strengthens the conclusion that late-passage immortalized cells have catastrophically lost early myogenic identity programming.

Strikingly, MYH8, MYH9, and MYH11 (myosin heavy chain isoforms) remained robustly expressed at the protein level despite transcriptomic alterations, demonstrating functional decoupling between early specification factors (MYOD1/MYOG) and terminal contractile machinery. This discordance likely reflects the exceptionally long protein half-life of sarcomeric myosin (estimated 3-4 hours for replacement in thick filaments, but overall protein turnover of ∼6 days), combined with potential post-transcriptional regulation through RNA-binding proteins that selectively stabilize or enhance translation of myosin transcripts. Studies by Ojima ^33^demonstrated that myosin molecules are continuously exchanged in thick filaments of cultured myotubes with a replacement half-life of 3-4 hours, yet overall myosin protein turnover (synthesis to degradation) requires ∼6 days, indicating remarkable protein stability once incorporated into sarcomeric structures. Furthermore, widespread mRNA-protein discordance has been documented across differentiation processes, with post-transcriptional mechanisms (selective translation, mRNA localization, RNA-binding protein regulation) enabling protein expression patterns divergent from transcriptional programs^33–36^.

Cell cycle regulators exhibit post-transcriptional complexity. CDKN1A (p21) protein showed dramatic upregulation (6.78-fold) despite mRNA suppression, exemplifying stress-responsive protein stabilization through deubiquitylation and reduced proteasomal degradation. p21 is normally unstable (20-60 minute half-life) but can be stabilized through deubiquitination by USP11, acetylation by Tip60, and complex formation that prevents ubiquitination. This protein stabilization likely represents cellular stress responses attempting to preserve growth checkpoints during immortalization^37,38^.

CDKN1B (p27) showed concordant suppression at both levels, indicating distinct regulation from p21. CCND1 (cyclin D1) exhibited mRNA downregulation but stable protein expression, consistent with protein stabilization through CDK4 complex formation. Studies demonstrate that cyclin D1 binding to CDK4 protects it from proteasomal degradation, explaining maintained protein despite endogenous gene suppression^39^.

In cell proliferation markers, CCNE1 (Cyclin E1) and MKI67 (Ki-67) showed stable expression between LPILM and PLM cells, indicating that immortalization extends cellular lifespan without inducing hyperproliferative phenotypes characteristic of oncogenic transformation. RB1 showed downregulation while PIK3R1 (PI3K regulatory subunit 1) showed upregulation, suggesting weakened G1/S checkpoints compensated by enhanced PI3K/AKT pathway responsiveness, a balanced adaptation supporting sustained proliferation while avoiding uncontrolled growth^40^.

BAX (pro-apoptotic) and BCL2 (anti-apoptotic) levels remained unchanged, demonstrating that LPILM cells retain normal apoptotic machinery balance despite immortalization. This finding provides crucial biosafety validation, confirming that CDK4/hTERT immortalization does not confer apoptosis resistance characteristic of oncogenic transformation, maintaining cellular quality control mechanisms essential for food safety applications^25^.

In addition, TP53 and ATM protein levels remained stable (ATM: 0.92-fold, log_2_FC = -0.11), concordant with transcriptomic findings showing no significant changes, validating that CDK4/hTERT bypasses rather than inactivates tumor suppressors, preserving genomic stability checkpoints.

#### 3.3.3 ECM remodeling: Definitive multi-omics validation of species-specific response

Proteomic analysis provided definitive validation of the extensive ECM remodeling identified transcriptomically, with perfect concordance between mRNA and protein changes demonstrating that structural ECM dismantling occurs coordinately at both regulatory and functional levels.

COL1A1 showed dramatic protein downregulation (0.27-fold, log_2_FC = -1.86), directly corroborating transcriptomic suppression. COL16A1 was completely absent in immortalized cells at the protein level (0-fold change), validating severe transcriptomic suppression. This 100% protein-mRNA concordance provides definitive evidence that immortalization triggers ECM structural dismantling at both transcriptional and translational levels in lamb muscle cells.

Additional structural proteins showed coordinated suppression: LRCH2 (leucine-rich repeats and calponin homology domain containing 2), STOM (stomatin), and PHF10 (PHD finger protein 10) were downregulated, indicating systematic remodeling of cytoskeletal organization and membrane architecture. SLC3A2 (CD98 heavy chain) demonstrated downregulation at both protein and mRNA levels (transcriptomic log2FC = -1.18 to -1.19), validating altered amino acid transport and integrin signaling.

#### 3.3.4 Metabolic enzyme upregulation: Functional validation of enhanced biosynthetic capacity

Proteomic analysis validated and extended transcriptomic predictions of metabolic reprogramming, with dramatic upregulation of enzymes supporting glycolysis, pH homeostasis, redox balance, and biosynthetic pathways.

Carbonic anhydrase isoforms CA2 and CA8 showed >10-fold protein upregulation, providing functional validation of transcriptomic predictions. Carbonic anhydrases catalyze reversible CO₂ hydration to bicarbonate and protons, critical for pH homeostasis under intense glycolysis characteristic of highly proliferative cells. The massive protein-level upregulation confirms that immortalized cells engage active pH buffering to maintain intracellular pH despite enhanced metabolic acid production.

GCLM (glutamate-cysteine ligase modifier subunit) showed >2-fold upregulation, validating enhanced glutathione biosynthesis predicted by transcriptomics. GCLM is the rate-limiting regulatory subunit for glutathione synthesis, and its protein-level elevation confirms functional enhancement of antioxidant capacity essential for managing oxidative stress in immortalized cells under prolonged culture^41,42^.

Nucleotide metabolism and biosynthetic pathways. NT5E (ecto-5’-nucleotidase, CD73) exhibited significant upregulation (>2-fold), concordant with transcriptomic findings, validating enhanced purine salvage and adenosine generation. H6PD (hexose-6-phosphate dehydrogenase) demonstrated >3-fold protein upregulation, confirming transcriptomic predictions and validating enhanced pentose phosphate pathway engagement for NADPH generation and ribose-5-phosphate biosynthesis essential for rapidly proliferating cells^43^.

#### 3.3.5 Cell adhesion and signaling molecules: Functional ECM interaction remodeling

LGALS3 (Galectin-3) showed >2-fold protein upregulation, concordant with transcriptomic findings, validating enhanced cell-cell and cell-matrix adhesion. Galectin-3 modulates cell adhesion, apoptosis resistance, and inflammatory responses, and its protein-level elevation confirms functional adaptation supporting immortalized cell survival despite ECM structural protein depletion^44–46^. STRA6 (stimulated by retinoic acid 6) demonstrated >3-fold protein upregulation, functioning as cell surface receptor for retinol-binding protein mediating vitamin A uptake. Recent studies highlight STRA6’s roles in muscle stem cell quiescence maintenance and mitochondrial function protection, and its dramatic upregulation suggests immortalized cells engage retinoid signaling pathways for metabolic adaptation and stress mitigation^47^. DACH1 (dachshund homolog 1) showed significant upregulation (>2-fold), functioning as transcriptional cofactor involved in organ development and cell fate specification. Its elevation may represent compensatory mechanisms attempting to maintain developmental regulatory capacity despite suppression of core myogenic transcription factors^48^.

#### 3.3.6 Growth factor signaling: Protein-level validation

IGFBP2 demonstrated dramatic protein downregulation (0.17-fold, log_2_FC = -2.53), showing perfect concordance with transcriptomic suppression, representing 100% multi-omics agreement. This definitive validation confirms fundamental IGF signaling alterations during immortalization, with IGFBP2 suppression potentially permitting sustained proliferation by reducing differentiation-promoting signals^49^.

#### 3.3.7 S100 protein family: Multi-omics validation of inflammatory activation

The S100 protein family showed coordinated upregulation validating transcriptomic inflammatory signatures. S100A4 demonstrated significant protein upregulation (1.71-fold, log_2_FC = 0.77), S100A16 showed stronger elevation (1.86-fold, log_2_FC = 0.90), and S100G exhibited dramatic increase (3.75-fold, log_2_FC = 1.91). These findings provide protein-level validation of inflammatory pathway activation.

S100 proteins function as calcium-binding DAMPs (damage-associated molecular patterns) activating inflammatory signaling and cellular stress responses. Their coordinated protein-level upregulation confirms that immortalization, despite its targeted mechanism avoiding oncogene-induced stress, still triggers functional inflammatory pathway activation in lamb muscle cells, a species-specific response with implications for long-term culture stability and potential immunogenicity in food applications^50^.

#### 3.3.8 Gene Ontology enrichment: Coordinated functional category alterations

Proteomic GO enrichment analysis revealed coordinated functional reorganization across biological processes (BP), cellular components (CC), and molecular functions (MF), validating transcriptomic predictions while providing functional confirmation at the protein level through complementary visualization approaches (Figure 6a, b).

**Figure 6.**
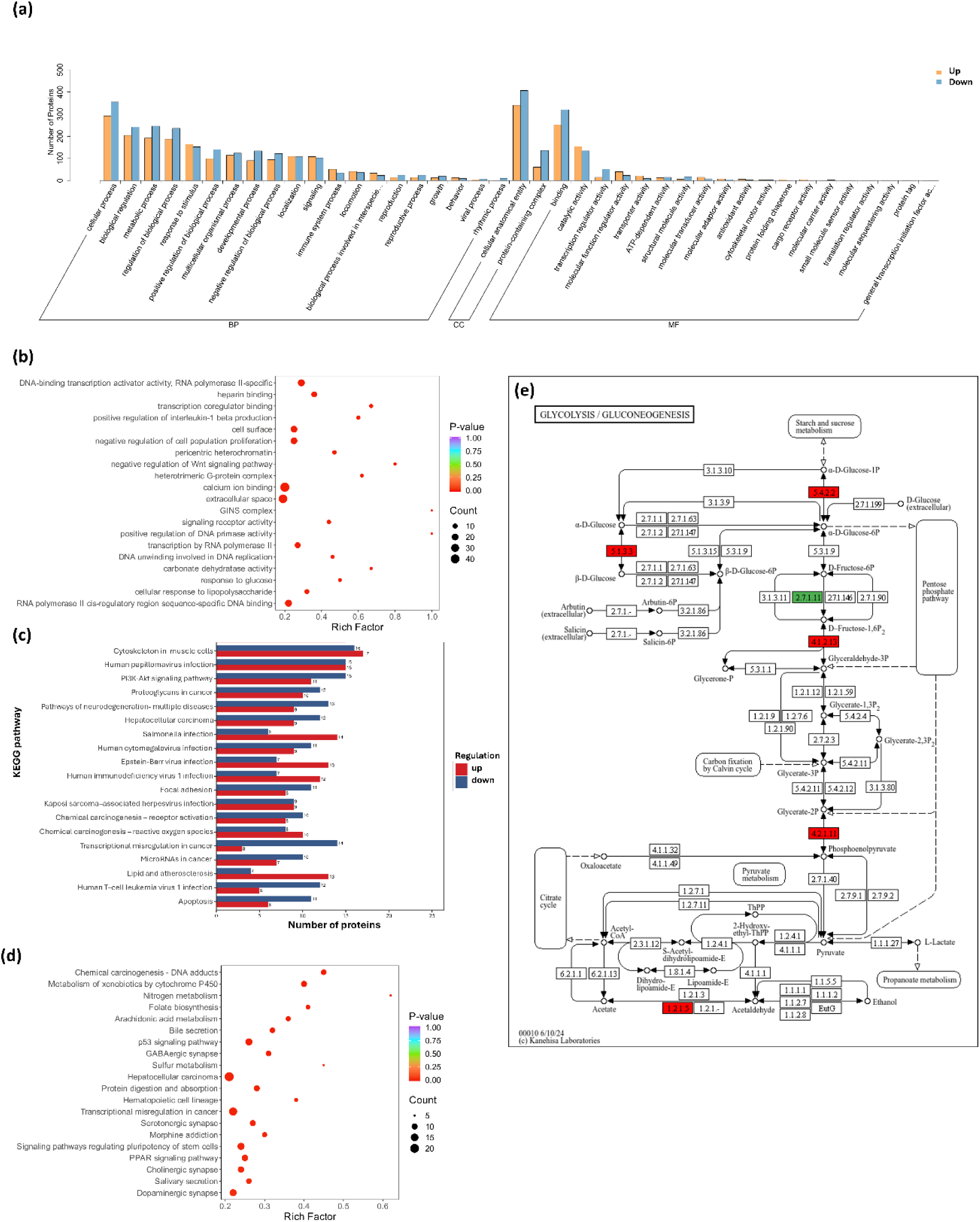
Comprehensive Gene Ontology and KEGG pathway enrichment analysis validates coordinated proteomic reorganization and metabolic reprogramming. (a) GO enrichment bar chart showing up-regulated (orange) and down-regulated (blue) protein counts across biological processes (BP), cellular components (CC), and molecular functions (MF). Prominent patterns include ECM/structural protein downregulation (blue bars exceeding orange in extracellular matrix organization, extracellular structure organization categories), balanced metabolic protein regulation (similar orange/blue bar heights in cellular component assembly, localization categories), and calcium ion binding enrichment, validating transcriptomic functional predictions at protein level. (b) GO enrichment bubble plot displaying significantly enriched GO terms. ECM categories (extracellular space, extracellular matrix organization) appear as larger orange bubbles confirming primary immortalization signature; metabolic and stress response categories prominently featured validate transcriptomic GO predictions at functional protein level; transcription-related terms show moderate significance confirming regulatory network suppression. (c) KEGG pathway bar chart showing protein counts for top enriched pathways with directional regulation (red = upregulated, blue = downregulated). “Cytoskeleton in muscle cells” shows balanced regulation (17 up, 16 down) confirming dynamic remodeling while preserving muscle functionality; “Pathways in cancer” shows predominant downregulation (24 down vs 15 up) supporting non-oncogenic profile; metabolic pathways (glycolysis, pentose phosphate) and stress response pathways (chemical carcinogenesis, drug metabolism) show enrichment confirming functional reprogramming. (d) KEGG pathway bubble plot showing significantly altered pathways with statistical significance representation. (e) Glycolysis/gluconeogenesis pathway diagram showing enzyme-level regulation (red boxes = upregulated, green boxes = downregulated, white boxes = unchanged), mechanistically validating transcriptomic metabolic predictions (22 metabolic genes in LPILM vs EPILM). Enhanced downstream glycolytic flux (red boxes clustering in fructose-1,6-bisphosphate → pyruvate segment: aldolase, enolase, phosphoglucomutase upregulated) with maintained upstream regulatory control (hexokinase, phosphofructokinase stable/downregulated at entry points) demonstrates coordinated metabolic adaptation supporting proliferation while avoiding uncontrolled glycolytic acceleration characteristic of oncogenic Warburg effect.

DNA-binding transcription activator activity showed coordinated suppression with 19 of 22 proteins downregulated, providing perfect functional validation of transcriptomic suppression of myogenic transcription factors (PAX3, MYOD1, MYOG). This protein-level confirmation demonstrates that transcriptional regulatory network collapse observed in RNA-seq translates to actual depletion of transcription factor proteins. The bubble plot representation in Figure 6b further illustrates this pattern with transcription-related GO terms showing moderate bubble sizes with high statistical significance, confirming systematic functional suppression

Cell surface and extracellular space categories showed enrichment, validating that transcriptomic ECM reorganization (identified as most significantly enriched biological process in LPILM vs PLM) translates to functional protein-level restructuring. The proteomic enrichment of ECM categories with predominant downregulation confirms systematic remodeling of cell-environment interactions. Despite structural protein loss, cells maintain viability through compensatory adhesion mechanisms (LGALS3 upregulation), demonstrating adaptive survival strategies during ECM dismantling.

Calcium ion binding (5.79% enrichment) and carbonate dehydratase activity showed prominent enrichment, functionally validating transcriptomic predictions of enhanced pH buffering (CA2/CA8 upregulation) and altered calcium homeostasis (46 genes in transcriptomic calcium ion binding category). Glutathione biosynthetic processes enrichment confirms GCLM-mediated redox adaptation predicted transcriptomically.

Negative regulation of cell population proliferation and Wnt signaling pathway categories (up to 2.81% enrichment) reflected preserved checkpoint control mechanisms, concordant with transcriptomic evidence maintaining non-oncogenic profiles and validating that immortalization does not induce uncontrolled hyperproliferation.

#### 3.3.9 KEGG pathway analysis: Balanced network reprogramming with non-oncogenic profile

Proteomic KEGG pathway analysis validated transcriptomic pathway predictions while revealing functional-level coordination through both quantitative distribution analysis (bar charts) and statistical significance assessment (bubble plots), providing comprehensive confirmation of pathway-level molecular reprogramming (Figure 6c-e).

Cancer-related pathways showed predominant protein downregulation (Figure 6c, 24 downregulated vs 15 upregulated proteins), providing functional validation of transcriptomic findings showing no oncogenic pathway activation. This balanced proteomic signature supports safety profile maintenance throughout extensive molecular reprogramming.

Cytoskeleton in muscle cells pathway displayed balanced protein changes (17 upregulated, 16 downregulated), functionally validating transcriptomic evidence of cytoskeletal pathway disruption (significant at protein level, marginally significant transcriptomically in early stage, highly significant in late stage). The balanced proteomic pattern confirms dynamic remodeling while preserving essential muscle structural features through selective protein retention (MYH8/9/11 stability).

Analysis on focal adhesion and metabolic pathways shows enhanced focal adhesion pathway proteins which support persistent cell expansion capacity, validating transcriptomic predictions of enhanced PI3K-Akt signaling and ECM-receptor interaction alterations. Glycolysis/gluconeogenesis pathway showed coordinated protein upregulation of phosphoglucomutase, aldose 1-epimerase, fructose-bisphosphate aldolase, and enolase (Figure 6e), with 6-phosphofructokinase downregulated, providing mechanistic protein-level confirmation of the metabolic reprogramming predicted transcriptomically and suggesting coordinated metabolic control with enhanced downstream flexibility.

#### 3.3.10 Evolutionary conservation analysis: KOG and InterPro domain validation

KOG (EuKaryotic Orthologous Groups) analysis reinforced proteomic findings with significant enrichment in extracellular structures (22 protein hits, representing 28% of all significant KOG categories), signal transduction mechanisms (130 hits, representing the largest category), and cytoskeletal organization. This distribution shows strong concordance with transcriptomic KOG analysis, demonstrating evolutionary conservation of affected processes and validating that immortalization-induced changes impact fundamental, evolutionarily conserved cellular systems at both mRNA and protein levels.

InterPro domain analysis revealed systematic protein-level downregulation of helix-loop-helix DNA-binding domains and homeobox domain superfamilies, directly matching transcriptomic suppression of early myogenic commitment programs and providing architectural-level validation of transcription factor network collapse. Conversely, carbonic anhydrase domains and metabolic enzyme domains showed protein-level enhancement, functionally validating coordinated metabolic adaptations supporting robust survival and proliferation predicted from transcriptomic metabolic pathway enrichment (22 metabolic genes in LPILM vs EPILM).

These data highlight the necessity of species-specific responses to immortalization, as documented in sheep cells^10^. Our multi-omics data further confirm that ovine muscle cells undergo more extensive molecular reprogramming compared to their human counterparts^5^, underscoring the need for species-optimized protocols.

### 3.4 Immunofluorescence Validation Reveals Post-Transcriptional Regulation of Myogenic Markers

To validate myogenic characteristics at the single-cell protein level, we performed immunofluorescence staining for PAX7, desmin, and myosin heavy chain (MHC) in PLM, EPILM, and LPILM cells (Figure 7). Immunofluorescence detected PAX7 protein in all three cell populations (Figure 7a), despite progressive transcriptomic suppression and no detection by mass spectrometry. Desmin and MHC remained positive despite reduced bulk omics levels (Figure 7b,c). Positive immunofluorescent staining despite low or undetected signal by mass spectrometry likely reflects a combination of spatial and biological factors. Proteins concentrated within the nucleus or other subcellular compartments are readily detected by spatial imaging but become diluted in total cell lysates, reducing detectability by mass spectrometry. Similarly, proteins expressed in rare subpopulations of cells may produce strong localized fluorescence yet contribute negligibly to the bulk proteomic signal. Immunofluorescence also benefits from the higher sensitivity of antibody-based detection, particularly for low-abundance targets that often fall below the dynamic range of conventional MS workflows. Furthermore, post-transcriptional mechanisms such as acetylation-mediated stabilization can preserve protein levels even when mRNA expression is suppressed, resulting in protein persistence despite transcript downregulation^51,52^

**Figure 7.**
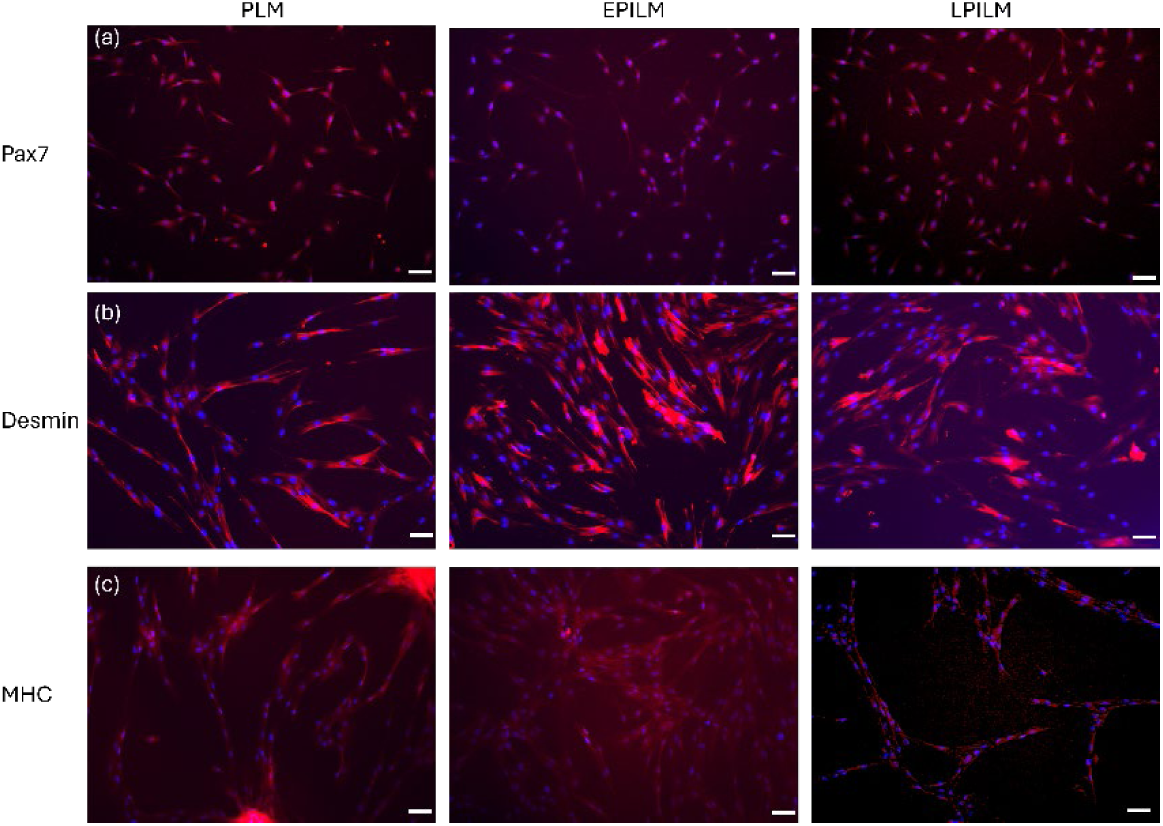
Immunofluorescence validation of myogenic markers across immortalization stages confirms preserved differentiation capacity. Representative immunofluorescence images of PLM , EPILM, and LPILM cells following 25 days of differentiation induction. (a) PAX7 immunostaining (red) with DAPI nuclear counterstain (blue) demonstrates sustained PAX7+ satellite cell populations across all passages despite dramatic transcriptomic suppression in immortalized cells. (b) Desmin immunostaining (red) with DAPI reveals maintained cytoskeletal intermediate filament networks across passages. Positive desmin signal despite reduced mRNA/protein levels in bulk omics reflects desmin’s extended protein half-life, incorporation into stable cytoskeletal structures, and high antibody detection sensitivity. (c) Myosin heavy chain (MHC) immunostaining (red) with DAPI following differentiation validates functional contractile machinery preservation across passages. Scale bars = 100 μm.

### 3.5 Implications of Ovine Satellite Cell Immortalization for Cultivated Meat Production

Our results demonstrate that CDK4/hTERT immortalized ovine satellite cells possess the essential attributes required for scalable and reliable cultivated meat production while revealing key biological and engineering challenges that must be addressed for commercialization. The long-term proliferative stability of these cells across more than 67 passages, coupled with consistent doubling times and cryostability, confirms that they can support continuous biomass generation without donor dependency, meeting one of the foremost requisites for cellular agriculture scalability. The sustained expression of fundamental myogenic proteins such as desmin and myosin heavy chain, and the preservation of differentiation potential—most notably in early-passage immortalized cells—validate these lines as a stable muscle cell source capable of recreating the texture and contractile protein content typical of natural meat. However, integrated transcriptomic–proteomic analysis revealed adaptive molecular reprogramming associated with immortalization that carries implications for both texture and product quality. The marked suppression of extracellular matrix components such as elastin and collagen indicates that cell-only constructs generated from these lines are likely to yield softer, less fibrous muscle tissue unless supplemented with ECM scaffolds or co-cultured with fibro-adipogenic cells to restore structural integrity. Upregulation of inflammatory S100 proteins and persistent carbonic anhydrase activity reflect stress-adaptive, glycolytic metabolic states, which may aid high-density culture resilience but could alter nutrient content or flavor precursors in the finished product. The metabolic shift toward enhanced glycolysis, while advantageous for rapid growth in bioreactors, also mirrors the Warburg-like phenotype seen in immortalized lines and necessitates careful monitoring to avoid metabolic imbalance or off-flavor compound accumulation. Importantly, the absence of transcriptomic activation in oncogenic pathways suggests that these cells remain biosafe for food applications, although additional functional tests—karyotyping, anchorage independence, and long-term tumorigenicity assays—would be mandatory for regulatory approval and consumer confidence. From an industrial perspective, the robust growth kinetics, serum-free adaptability, and predictable differentiation profile make these ovine cells an excellent production chassis for lamb-specific cultivated meat products; nevertheless, improving ECM deposition, controllable lipid incorporation, and sensory mimicry remains the next stage of optimization. Altogether, the findings situate these immortalized ovine muscle cells as a viable and tunable biofactory platform for cultured meat production, establishing foundational insights into how immortalization reshapes cell physiology, guides media formulation, and defines both technical feasibility and sensory potential of next-generation cellular meats.

## Conclusion

This study provides the first comprehensive multi-omics characterization of CDK4/hTERT immortalized ovine satellite cells, revealing fundamental insights into cellular reprogramming relevant to cultivated meat applications. Our integrated transcriptomic-proteomic approach demonstrates that immortalization triggers progressive molecular evolution involving coordinated transcriptional and post-transcriptional adaptations that intensify with passage progression. Despite extensive molecular reprogramming affecting >1,000 genes and 960 proteins, immortalized cells retain essential characteristics for cultivated meat applications: preserved contractile proteins, maintained apoptotic machinery, non-oncogenic pathway profiles, and stable proliferation through 64 passages. Critically, some transcriptome-proteome discordance highlights the importance of multi-omics approaches for comprehensive cell line characterization.

The established ovine satellite cell line addresses scalability challenges in lamb meat production while providing a robust platform for regulatory assessment. Our systematic multi-omics framework establishes new standards for immortalized cell line characterization that balance the scalability requirements of commercial cultivated meat production with the biological authenticity necessary for safe, high-quality products.

## Acknowledgments

This research was financially supported by the Agriculture and Food Research Initiative (AFRI) Sustainable Agricultural Systems program, grant no. 2021-699012-35978 from the USDA National Institute of Food and Agriculture and Texas A&M AgriLife Research.

## Data Availability Statement

Data will be made available on request.

